# Lysophosphatidylcholine acyltransferase 2 (LPCAT2) Regulates The Expression of Macrophage Phenotype Markers in RAW264.7 Cells

**DOI:** 10.1101/2021.11.25.469956

**Authors:** Victory Ibigo Poloamina, Hanaa Alrammah, Wondwossen Abate, Neil Avent, Gyorgy Fejer, Simon K. Jackson

## Abstract

Macrophages are key antigen presenting cells that also secrete cytokines during inflammation. They can be polarised to M1 or M2 phenotypes. Molecules such as CD206 and inducible Nitric Oxide Synthase are considered macrophage phenotype markers because they are highly expressed in either M1 or M2 macrophages. LPCAT2 is a phospholipid modifying enzyme that influences inflammatory responses in macrophages. However, how LPCAT2 influences inflammation is not fully understood.

In this study, we have used genetic technology to study the influence of LPCAT2 on macrophage phenotype markers.

Our results show for the first time that overexpression of LPCAT2 promotes the expression of M1 macrophage phenotype markers, and attenuates the expression of M2 macrophage markers.

## 2 Introduction

Macrophages are primary effector cells of the innate immune system, that differentiate from monocytes. They belong to a group of phagocytic cells that stem from Myeloid progenitors. These cells have a professional function of antigen-presentation and are characterised with the ability to secrete cytokines that attract other immune cells and initiate inflammation in response to pathogens at the site of infection [1]. Macrophages can be polarized into classically activated or alternatively activated phenotypes. Classically activated macrophages are also known as M1 or M1-like macrophages because they preferentially promote Th1 (T Helper 1) cell responses; thus they are pro-inflammatory and release anti-microbial molecules such as nitric oxide [1, 2]. On the other hand, alternatively activated macrophages preferentially promote Th2 (T Helper 2) cell responses, therefore they are often referred to as M2 or M2-like macrophages. M2 macrophages mainly phagocytose pathogens, and often have a higher expression of mannose and scavenger receptors compare to M1 macrophages. They also promote cell proliferation and tissue repair [3, 4].

Certain molecules have been classed as markers of macrophage polarization due to their consistently high expression in either phenotypes of macrophages. In M1 macrophages, molecules such as inducible Nitrogen Oxide Synthase (iNOS), CXCL10, Type I Interferons, Tumour Necrosis Factor Alpha (TNF⍰). Whereas, in M2 macrophages, molecular markers include CD206, CD36, Interleukin 10. M2 macrophages also exhibit increased fatty acid oxidation, and adipogenesis, and a higher expression of molecules involved in fatty acid metabolism such as carnitine palmitoyltransferase, (PPAR⍰) [5].

LPCAT2 (Lysophosphatidylcholine Acyltransferase) also known as LysoPAF acetyltransferase localises in macrophage membranes and lipid droplets [6, 7]. It preferentially uses arachidonic-CoA as a donor for the acylation of fatty acids, thereby producing several varieties of arachidonic acid. Arachidonic acid is a key precursor for the synthesis of lipid mediators such as prostaglandins, leukotrienes, and lipoxins. Therefore, LPCAT2 influences inflammation by regulating the synthesis of fatty acids and lipid mediators such as PAF, Leukotriene B4, and Prostaglandin E2, and the expression of proteins involved in macrophage inflammatory responses to lipopolysaccharide (LPS) [8, 9]. Further research is needed to understand the mechanism of LPCAT2 involvement in macrophage inflammatory responses. In this paper, we propose that LPCAT2 regulates macrophage inflammatory responses by influencing the expression of molecular markers of macrophage polarization.

## 3 Materials and Methods

Chemicals reagents used to prepare buffers and BCA Assay kit were purchased from Sigma Aldrich, UK and Fisher Scientific, UK. Buffers used include RIPA buffer, Phosphate-buffered saline (PBS), Tris-buffered saline (TBS), Blocking Buffer, Cell lysis buffer, elution buffer, SDS sample buffer, and ECL detection reagent [23]. PolyPlus Lipofectamine 2000 was purchased from Source Bioscience, UK. Opti-MEM, Power SYBR Green, RNa to cDNA kit, and gel casting materials were purchased from Life Technologies, UK. Antibodies, Protein A/G agarose gel beads, and protein ladders were obtained from Santa Cruz Biotechnology, UK and Cell Signalling Technologies, UK. DMEM culture medium and other cell culture materials were purchased from Lonza, UK. PCR primers were designed with Primer3 Plus Bioinformatics Softwaew and NCBI BLAST, and purchased from Eurofins Genomics.

### 3.1 Culturing of RAW264.7 Cell Line

RAW264.7 cell line was obtained from the European Collection of Cell Cultures (ECACC) through Public Health England, UK. RAW264.7 macrophages were maintained in Dulbecco’s Modified Eagle Medium (DMEM) [Lonza, BE12-914F] supplemented with 10%(v/v) Foetal Bovine Serum (FBS) [Labtech.com, BS-110] and 1%(v/v) 0.2M L-Glutamine [BE17-605E], and incubated at 37°C, 5% CO_2_. Lipopolysaccharide (*E. coli* O111:B4) [Sigma-Aldrich, L2630] was resuspended in LAL reagent water (<0.005EU/ml endotoxin levels) [Lonza, W50-640]. Cell culture medium was used to dilute all ligands to the needed concentration before stimulation.

### 3.2 Expansion and Purification of Plasmids

Kanamycin-resistant TrueORF Gold plasmid was expanded by transformation of 1⍰g/ml pCMV6 plasmid vector [4.9kb; OriGene, PS100001] with or without murine LPCAT2 recombinant DNA [Open Reading Frame; Forward 5–CAG ACT GTT ACG GGCTTT GCA - 3; Reverse 5– ACC TGA TGT CGC TCG CTTTT - 3] in 10⍰l Alpha-Gold competent *E. coli* cells [Insight Biotechnology, UK] using GenElute™ plasmid MiniPrep Kit [Sigma-Aldrich, PLN10].

#### 3.3 Transfection of RAW264.7 Cells with Plasmid Containing LPCAT2 Insert

2.5μg of pCMV6 plasmid vector with or without murine LPCAT2 recombinant DNA was transfected into RAW264.7 macrophages using Lipofectamine 2000 transfection reagent [PolyPlus transfection, 114-07]. Macrophages were cultured in DMEM 24 hours before gene silencing, then in Opti-MEM (Reduced Serum Medium) for 24 hours during gene silencing and incubated at 37°C, 5% CO_2_. Prior to cell stimulation or any further experiments, Opti-MEM (Reduced Serum Medium) was replaced with DMEM cell culture medium for 24 hours and incubated at 37°C, 5% CO_2_. Transfection mixture was prepared using Opti-MEM.

### 3.4 Reverse Transcription and Real Time Quantitative PCR (RT-qPCR)

The reaction mastermix contained 37%(v/v) nuclease-free, 230nM of target primers (a mixture of both forward and reverse primers), 60%(v/v) Power SYBR Green, and 3μl of ≥ 100ng/μl cDNA. The reaction was initiated at 95°C for 10 minutes, then up to 40 repeated cycles of denaturing (15 seconds, 95°C), annealing, and extension (60 seconds, 60°C). GAPDH was used as an endogenous control. GAPDH [forward sequence, 5–CCT CGT CCC GTA GAC AAA ATG; reverse sequence, 5– TCT CCA CTT TGC CAC TGC AA]; LPCAT2 [forward sequence, 5–TGG AAG CGC CCA TTT TTG TC; reverse sequence, 5–GGG ATG AAG GCT CTG CCA AC]; iNOS [forward sequence, 5 – CGC CTT CAA CAC CAA GGT TG; reverse sequence, 5 – TCA GAG TCT GCC CAT TGC TG]; COX2 [forward sequence, 5–CCA CAG TCA AAG ACA CTC AGG TAG A; reverse sequence, 5– CCA GGC ACC AGA CCA AAG AC]; PPAR⍰ [forward sequence, 5–CCA CTC GCA TTC GTT TGA CA; reverse sequence, 5 – TCG CTC AGC TCT TCC GAA GTG]; CD206 [forward sequence, 5– AAA TGG AGC CGT CTG TGC AT; reverse sequence, 5 – AAG TGC AAT GGA CAA AAT CCAA].

### 3.5 Immunoblotting

Equal amounts of whole cell lysates and eluates were separated on pre-cast SDS-PAGE gels and blotted on to a PVDF membrane using a blot module. The blots were blocked with 0.1% bovine serum albumin in PBS-0.1% Tween 20, probed with primary and then HRP-conjugated secondary antibodies. The target proteins separated on the PVDF membrane were detected and analysed using ImageJ.

### 3.6 Data Analysis

Statistical Analysis was carried out in R Statistical Programming Software and graphs were plotted using ggplot2 package. All independent experiments were repeated at least 3 times. Data represent Mean ± Standard Error of Mean. Paired T-Test with Dunnetts T3 multiple comparison test were used for statistical analysis.

All statistical tests were significant at 95% confidence interval, p ≤ 0.05.

## 4 Results

### 4.1 Cloning of RAW264.7 Cells Over-expressing LPCAT2

LPCAT2 was cloned in *E. coli* and over-expressed in RAW264.7 cells using vector plasmid carrying a neomycin-resistant gene. After 3 days, RAW264.7 cells transfected with empty vector pasmid (pCMV6) and treated with 700μg/ml of geneticin, however, RAW264.7 cells transfected with neomycin-resistant vector plasmid over-expressing LPCAT2 survived for up to 7 days [data not shown]. Thus, cells not over-expressing LPCAT2 were killed off, allowing for cloning of cells over-expressing LPCAT2.

### 4.2 Expression of LPCAT2 in RAW264.7 Cell Transfected with LPCAT2 siRNA or Vector Plasmid Over-Expressing LPCAT2

The gene and protein expression of LPCAT2 were analysed to confirm the over-expression of LPCAT2. Figure 1A shows that LPCAT2 gene is significantly higher in cells over-expressing LPCAT2 (4.18 ± 1.1, p = 0.05), and LPCAT2 protein increased (6-fold) in cells over-expressing LPCAT2 (Figure 1B&C).

**Figure 1:**
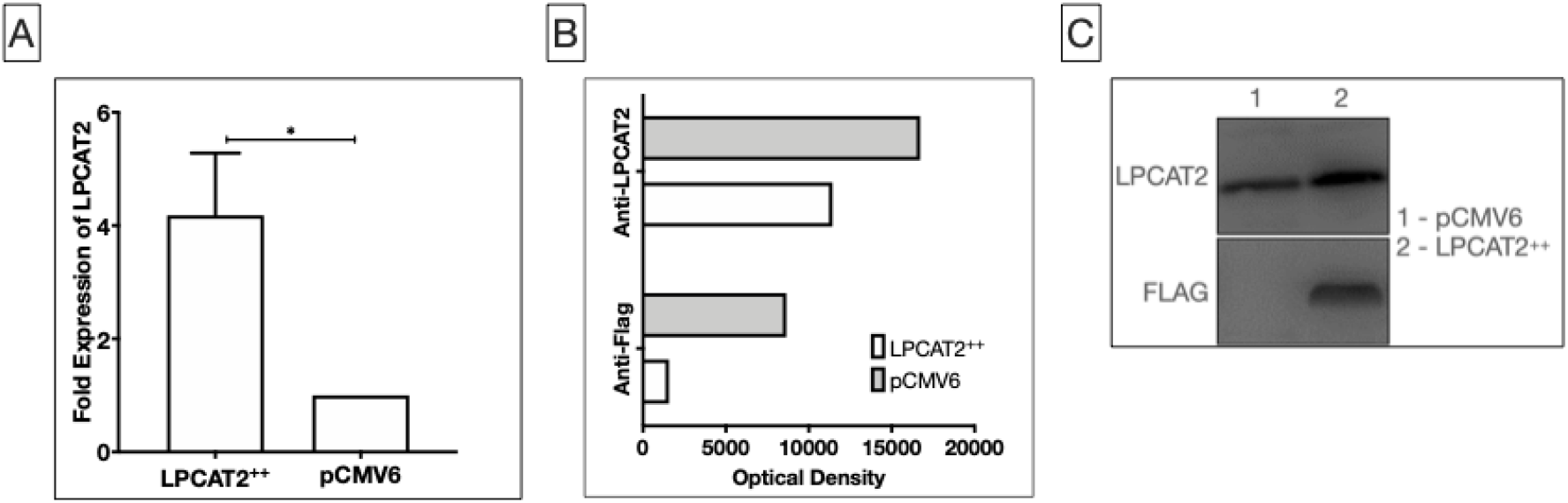
Over-expression of LPCAT2 in RAW264.7 cells caused ≥70 increase in LPCAT2 gene expression **(A)** and ≥50% increase in LPCAT2 protein expression **(B & C)**. pCMV6 is an empty vector plasmid, whereas LPCAT2 indicates cells over-expressing LPCAT2. *p≤0.05. Data represents mean of at least 3 independent experiments (n≥3) ± standard error (A), and optical densities (B) of western blots (C).

### 4.3 LPCAT2 Regulates the Expression of CD206, Inducible Nitric Oxide Synthase (iNOS), PPARγ, and Cyclo-oxygenase 2 (COX2)

Here, the effect of LPCAT2 over-expression on the gene expression of these macrophage phenotype markers was examined. Figure 2 shows that in RAW264.7 cells over-expressing LPCAT2, there was a significant increase in iNOS (281.43 ± 13.33, p < 0.0001) and COX2 (2659.39 ± 253.09, p = 0.0005) after treating the cells with 100ng/ml LPS. On the other hand, LPS significantly downregulated the gene expression of CD206 (0.89 ± 0.07, p = 0.03) and PPARγ (0.4 ± 0.004, p = 0.0009), and over-expression of LPCAT2 further decreased the gene expression of CD206 (0.7 ± 0.07, p = 0.1) and PPARγ (0.16 ± 0.003, p = 0.3). In non-treated cells, LPCAT2 over-expression significantly increased the gene expression of CD206 (3.22 ± 0.38, p = 0.0012) and iNOS (44.48 ± 8.46, p = 0.0021), but significantly decreased the gene expression of PPARγ (0.19 ± 0.03, p < 0.0001).

**Figure 2:**
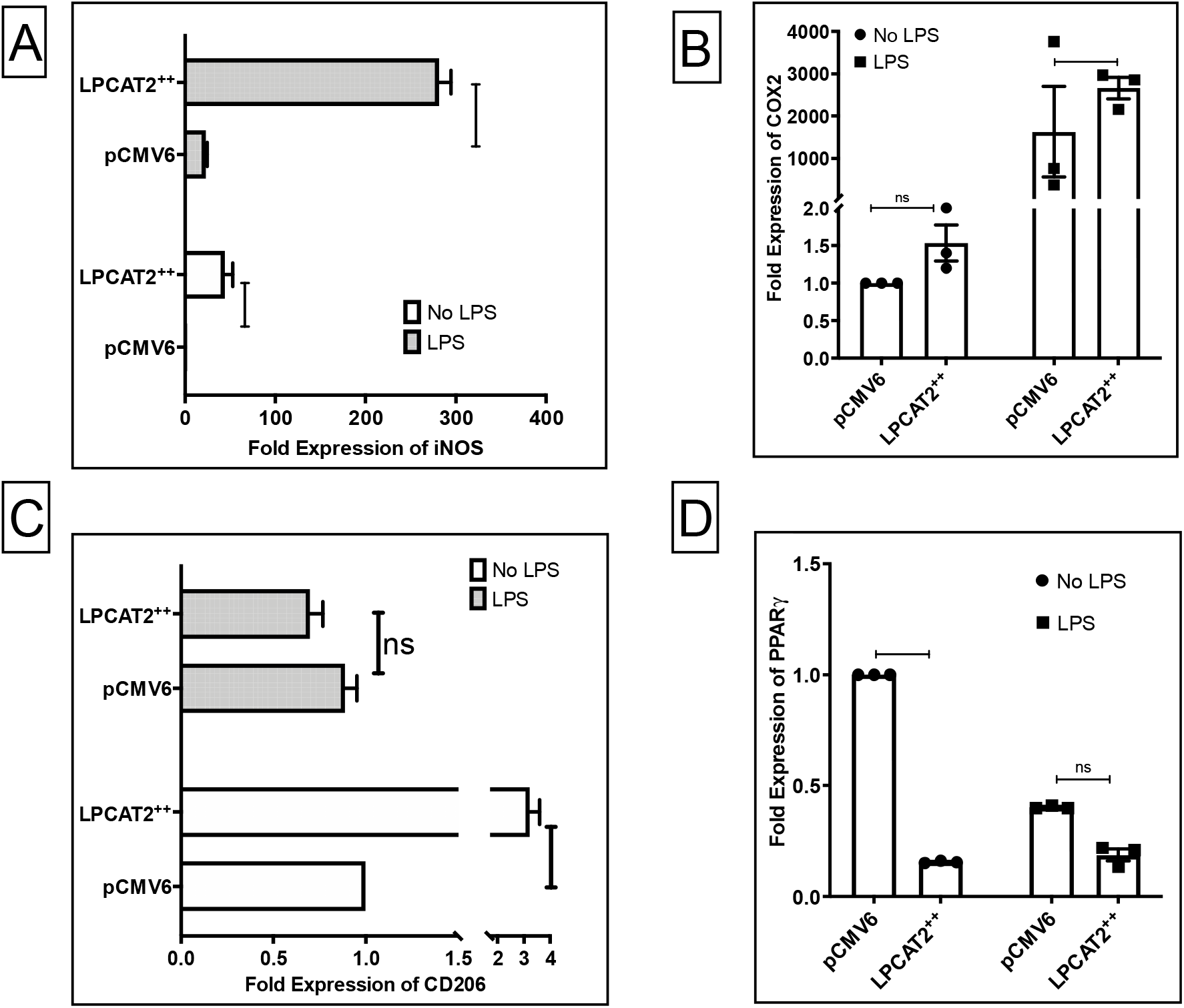
Over-expression of LPCAT2 increases the gene expression of inducible Nitric Oxide Synthase, iNOS **(A)** and Cyclo-oxygenase 2, COX2 **(B)**. On the other hand, over-expression of LPCAT2 results in a significant decrease in the gene expression of CD206 **(C)** and peroxisome proliferator-activated receptor gamma, PPARγ **(D)**. TLR4 ligand is 100ng/ml *E. coli* O111:B4 lipopolysaccharide. pCMV6 represents RAW264.7 cells transfected with empty vector plasmid and LPCAT2++ represents RAW264.7 cells over-expressing LPCAT2. *p≤0.05. Data represents the mean of 3 independent experiments (n=3) ± standard error.

### 4.4 LPCAT2 Expression is Significantly Increased in Primed RAW264.7 Cells

Figure 3 shows that priming RAW264.7 cells with 25ng/ml IFNγ for 24 hours and treating the cells for 2 hours with TLR4 ligand, LPS, significantly increases the gene expression of LPCAT2 (2.71 ± 0.63, p = 0.03). Likewise, priming RAW264.7 cells with a PPARγ antagonist, T0070907, and then treating the cells for 6 hours with LPS results in a dose-dependent increase in LPCAT2 gene expression (3.37 ± 0.84, p = 0.03; 5.38 ± 1.45, p = 0.002; 9.77 ± 1.72, p = 0.02) as seen in Figure 3B.

**Figure 3:**
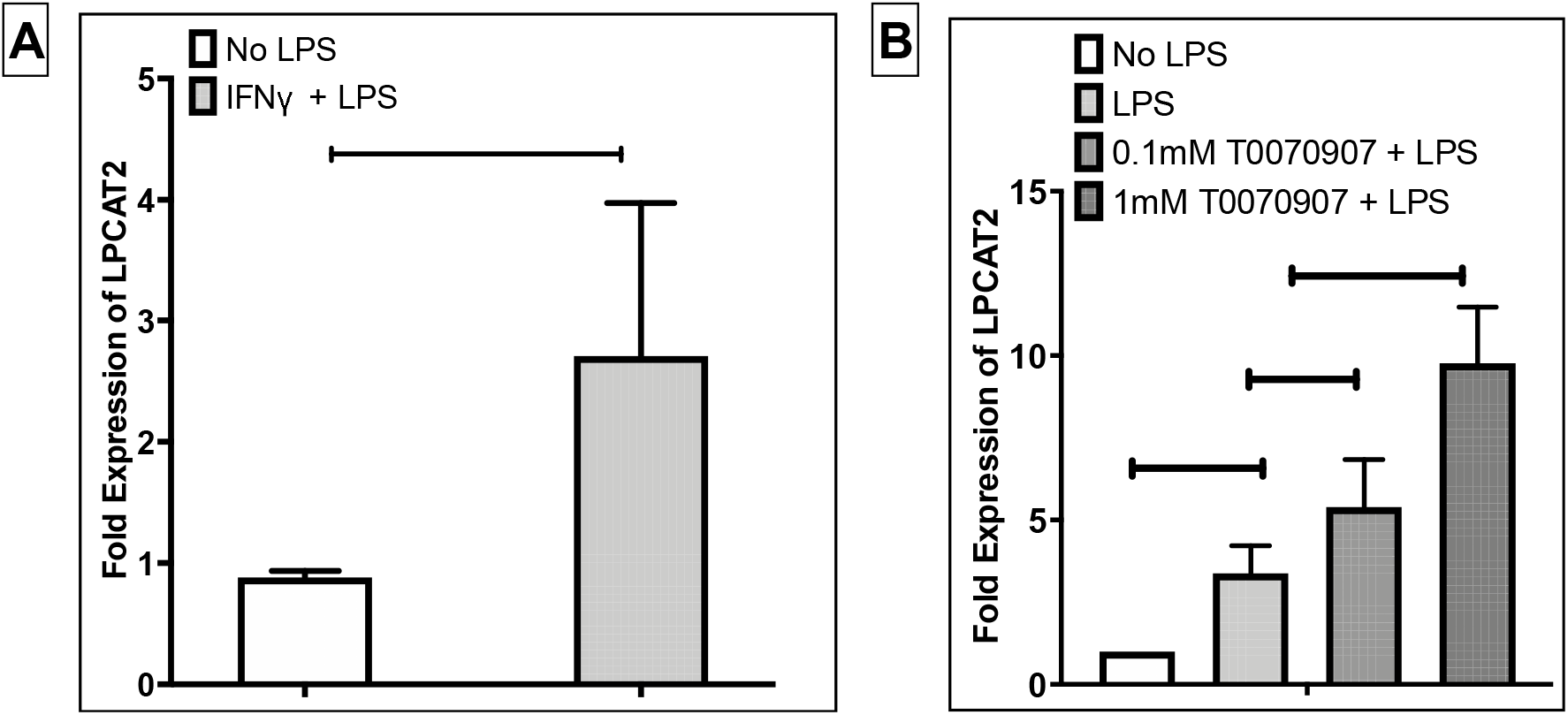
LPCAT2 gene expression is significantly increased in primed RAW264.7 cells. RAW264.7 cells were primed with either 25ng/ml of Interferon gamma (IFNγ) or varying concentrations of PPARγ antagonist (T0070907) and then stimulated with either 100ng/ml **(A)** or 1000ng/ml **(B)** of TLR4 ligand (*E. coli* O111:B4 lipopolysaccharide). *p≤0.05. Data represents mean of 3 independent experiments (n=3) ± standard error.

## 5 Discussion

COX2 and iNOS [5, 10] are common markers of M1 macrophages, and CD206 [11] and PPARγ [12] are markers of M2 macrophage. PPARγ receptor and TLR4 are some of the key receptors responsible for regulating macrophage activation to either M1 or M2 phenotypes [13]. During inflammation there is an increased release of chemokines and cytokines which can be regulated by various factors including Nitric oxide and prostaglandins [14]. COX2 synthesises prostaglandins by breaking down arachidonic acids – a catabolic process [15]. On the other hand, PPAR⍰ mainly store fatty acids as triglycerides in adipocytes – an anabolic process [16]. Anabolic processes are increased in M2 macrophages, whereas catabolic processes are increased in M1 macrophages [5]. Therefore, our results imply that LPCAT2 promotes catabolism and synthesis of lipid mediators, while inhibiting anabolism or adipogenesis (Figures 2 & 3). The inflammatory influence of LPCAT2 stems from its anabolic activity which builds up inflammatory mediator – PAF, or Arachidonic acid – a precursor for lipid mediators.

Pro-inflammatory conditions induce an increased expression of LPCAT2 as seen in Figure 3, implying a possible increase in its enzymatic activity which would lead to higher amounts of arachidonic acid [18]. Accumulation of arachidonic acid increases the expression and activity of COX2 [19, 20]. This is in line with our results which shows that LPCAT2 over-expression induces an increased expression of COX2 after stimulation with LPS. Furthermore, Over-expression of LPCAT2 significantly increased iNOS expression, both in LPS-stimulated and nonstimulated macrophages. The synthesis of nitric oxide by nitric oxide synthases and its release are hallmarks of inflammation [8], and inducible nitric oxide synthase (iNOS also known as NOS2), is classed as an M1 marker because of its consistently high expression in pro-inflammatory macrophages [5].

Cluster of Differentiation molecules are widely used to identify immune cell phenotypes and function, CD206 belongs to this class of molecules [23]. CD206 expression was not induced in LPS-stimulated macrophages over-expressing). CD206 is expressed in both M1 and M2 macrophages. CD206 (a mannose receptor) is commonly known as an M2 macrophage marker because it is highly expressed in M2 macrophages, and is induced predominantly by interleukin 4 (which induces M2 macrophage polarization) [5, 22]. A scientific publication claimed that there is no difference in CD206 mRNA expression in M1 and M2 macrophage phenotypes, however, the significant difference may have been masked by the variability of CD206 mRNA expression [2].

In conclusion, it is evident from the results that the expression of some macrophage phenotype markers are dependent on LPCAT2. Moreover, LPCAT2 promotes the expression of M1 markers while attenuating the expression of M2 macrophage markers. Therefore, this paper is further evidence that LPCAT2 is pro-inflammatory and regulates inflammation by inducing a higher expression of M1 markers.

